# Human amygdala volumetric patterns convergently evolved in cooperatively breeding and domesticated species

**DOI:** 10.1101/2023.05.30.542860

**Authors:** Paola Cerrito, Judith M. Burkart

## Abstract

**Purpose:** The amygdala is a hub in brain networks that supports social life and fear processing. Compared to other apes, humans have a relatively larger lateral nucleus of the amygdala, which is consistent with both the self-domestication and the cooperative breeding hypothesis of human evolution.

**Methods:** Here, we take a comparative approach to the evolutionary origin of the relatively larger lateral amygdala nucleus in humans. We carry out phylogenetic analysis on a sample of 17 mammalian species for which we acquired single amygdala nuclei volumetric data.

**Results:** Our results indicate that there has been convergent evolution toward larger lateral amygdala nuclei in both domesticated and cooperatively breeding mammals.

**Conclusion:** These results suggest that changes in processing fearful stimuli to reduce fear-induced aggression, which are necessary for domesticated and cooperatively breeding species alike, tap into the same neurobiological proximate mechanism. However, humans show not only changes in processing fearful stimuli but also in prosociality. Since cooperative breeding, but not domestication, is also associated with prosociality, a prominent role of the former during human evolution is more parsimonious, whereas self-domestication may have been involved as an additional stepping stone.

## 1. Introduction

The amygdala has been described as a hub in brain networks that support social life (Bickart et al., 2014; Wu & Hong, 2022). In humans, compared to other apes, the lateral nucleus of the amygdala is relatively larger (Barger et al., 2007) and contains more neurons (Barger et al., 2012). Recent work has demonstrated that in humans, reduced amygdala volume correlates with a reduction in the ability to both feel fear and recognize fear in others, and with an increased prevalence of psychopathic traits (Marsh et al., 2008). Similarly, extreme altruists are characterized by a heightened ability to recognize fear, and by a larger and more active amygdala (Marsh et al., 2014). These neuroanatomical findings appear to be well in line with the self-domestication (SD) hypothesis of human evolution (Hare, 2017), which argues for selection against fear-induced aggression toward strangers, given that the amygdala and particularly the lateral nucleus, play a critical role in fear processing.

However, this relative increase in lateral amygdala nucleus volume in humans compared to other apes, is also consistent with the cooperative breeding model (CB) of human evolution. The CB model (Hrdy & Burkart, 2020) posits that infant care by non-mothers (allomaternal care) became essential for infant survival during the Pleisotcene. Indeed, our life history pattern, which includes short interbirth intervals without compromising infant size and/or brain size, depends, energetically, on the availability of helpers in raising offspring. Importantly, this reliance on allomaternal care requires that immatures overcome the fear of being close to conspecifics who are not their mothers, even if these conspecifics are in the possession of valuable food sources in the case of provisioning. Moreover, mothers need to overcome the fear of others approaching, handling, and carrying their infants. These two different types of fear and consequent aggressive responses (affective and defensive) make use of the same underlying brain networks, as verified across several species of mammals (Panksepp, 2004).

In sum, both have to overcome fear, tolerate close proximity and eventually elicit care from others: domesticated species from humans, and cooperatively breeding ones from conspecifics. Both thus had to find ways to deal with these potentially fear-eliciting situations. We therefore predict similar neuroanatomical adaptations of the amygdala nuclei in both domesticated and cooperatively breeding species. We take a comparative approach to test whether the amygdala volumetric patterns observed in humans are more likely the result of SD or CB. Specifically, we hypothesize that a relatively larger lateral nucleus, which is the sensory interface in fear conditioning (LeDoux et al., 1990), allowed for the evolution of a heightened sensitivity in the recognition of others’ fear, expressed also, especially in cooperatively breeding species, via facial expressions (Cerrito & DeCasien, 2021). In humans, this increased ability to recognize others’ fear is thought to trigger a violence inhibition mechanism (Blair, 1995) which in turn allows for adults to care for infants that are not their own (Marsh, 2019).

To test our hypothesis, we combine neuroanatomical data with detailed data on infant care and domestication status and use phylogenetic methods to assess the correlation between relative volumes of different amygdala nuclei, domestication and CB and. Specifically, we test whether convergent evolution resulted in similar neuroanatomical adaptations in the amygdala of domesticated and cooperatively breeding species, with both groups having a larger than expected lateral nucleus than species who are neither domesticated nor cooperative breeders.

## 2. Materials and Methods

We acquired volumetric data of 27 adult individuals of 17 mammalian species (of which three domesticated, and three cooperative breeders) for the following regions: amygdaloid complex (AC), basolateral nucleus (BLA) and its lateral (L), basal (B) and accessory basal (AB) nuclei (the latter three add up to form the BLA). All volumetric data were obtained from the literature (Barger et al., 2007; Carlo et al., 2010; Chareyron et al., 2012; Równiak et al., 2022). When both left and right hemispheric values are reported, the mean was taken. Conversely, in some cases (Chareyron et al., 2011, 2012) the right or left hemisphere was randomly chosen after ascertaining the absence of lateralization (p=0.757) and yet in others (Carlo et al., 2010) only the left is reported. All behavioral data were taken from published literature (Bird et al., 2019; Cerrito & Spear, 2022; Dorning & Harris, 2019; Isler & van Schaik, 2012; Mauget, 1981; Nowak & Walker, 1999; Schweinfurth, 2020; Stone, 1995). For a definition of the measures of alloparental care see Isler and van Schaik (2012).

We created a binary variable aimed at expressing whether a species was a cooperative breeder. To do so we first created a compound “allocare_sum” variable as the sum of all forms of alloparental care observed in each species: male provisioning of infants (continuous), provisioning of infants by other group members (continuous), carrying of infants by males (continuous), carrying of infants by others (continuous), allonursing (continuous), protection by male (continuous), other forms of infant care such as thermoregulation, babysitting and pup retrieval (continuous). We then made this continuous variably binary (non-cooperative breeder = 0 to 2.64; cooperative breeder = 2.64 to 5.2) using k-means clustering implemented using the “discretize” function in the R package “arules” (Hornik et al., 2005). Finally, we made a compound binary variable, called “elicit_care”, which takes the value of 1 for species that are either domesticated or cooperative breeders, or the value of 0 for those that are neither. Given that in some species there isn’t infant carrying at all (nor maternal nor allomaternal), we also created an “allocare_sum_NoCarry” variable which represents the sum of all alloparental care behaviors, except infant carrying. The breaks used to discretize this variable are: non-cooperative breeder = 0 to 2.36; cooperative breeder = 2.36 to 4.5. The species resulting as cooperative breeders are the same three as in the “allocare_sum” variable, and therefore the value of elicit_care for each species does not change whether one selects the “allocare_sum” or the “allocare_sum_NoCarry” value. The final and complete dataset is provided in Data S1. Volume is expressed in cubic centimetres (cm^3^), weight in grams (g) and time in days (d).

We carried out all statistical analyses using R version 4.0.1 (R Core Team, 2020). We used phylogenetic generalized least squares (PGLS) regressions to assess the correlation between the variable “elicit_care” (domestication or cooperative breeding) and relative neuroanatomical data. To compare the data across species with a large range of variation in amygdala size, we regressed B, L and AB against BLA and against the entire AC. We used the additive effect of “elicit_care” to test for significant differences between intercepts. We assessed the correlation between infant care measures on one side and relative neuroanatomical data on the other using phylogenetic generalized least squares (PGLS) regressions (Grafen, 1989). The models used a recently updated species-level mammals phylogeny (Upham et al., 2019). We used the maximum clade credibility tree calibrated using node dates and an exponential prior. To account for the high degree of skew, we log-transformed (in base *e*) all the volumetric, weight and time variables.

We estimated λ for the PGLS models independently for each regression using maximum likelihood. We carried out the PGLS analyses using the “pgls” function in the R package “caper” (Orme et al., 2013). For each significant model we also report the effect size, expressed as percentage. To compare the effect size between different explanatory variables, we scaled the intercept difference to reflect the maximum value that the variable can take, so the intercept difference is multiplied by that value and then the difference in the natural log of the effect sizes is exponentiated to convert back to the original ratio of the effect sizes. We then proceeded to compute the AIC (Akaike, 1973) for each of the significant models and compared models with the same response variable. The complete list of models, their estimates and the results of the model selection using AIC are available in Table 1.

**Table 1.**
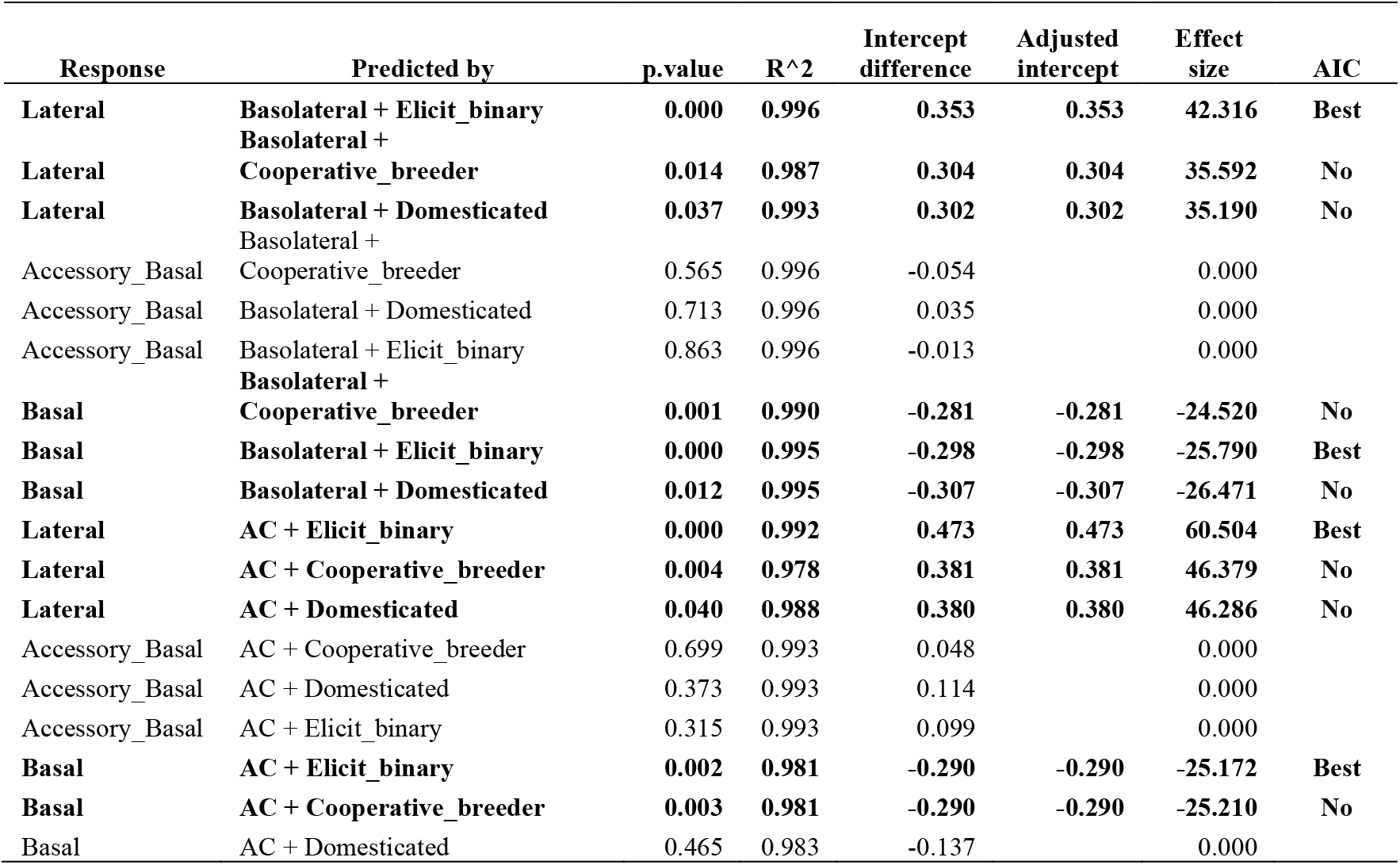
Estimates of the models in which each nucleus is regressed against the basolateral one or the entire amygdaloid complex (AC), plus the additive term tested. For each model: response variable, explanatory variable plus additive term, p-value, R^2^, intercept difference, adjusted intercept difference (see methods), and effect size (in %).’Best’ indicates that the model has the lowest AIC for that response variable and no other model is within 2; ‘Tied’ indicates that the model either has the lowest AIC but there are others within 2, or that its AIC is within 2 of the lowest; ‘No’ indicates that the AIC score is >2 from the lowest AIC score for that response variable.

As the results were significant, we proceeded to conduct post-hoc tests to assess which of the constituting variables of “elicit_care” (i.e. domestication and cooperative breeding) were, individually, correlated with significant nuclei volumetric differences.

For each continuous variable we measured the phylogenetic signal using Blomberg’s K (Blomberg et al., 2003) implemented by the “phylosig” function in the R package “phytools” (Revell, 2012). To test significance, we used 100,000 iterations. For discrete variables we followed previously published methods (Cerrito & Spear, 2022) and estimated the phylogenetic signal using δ (Borges et al., 2019), and ran 10,000 iterations to test for significance. The value of K/ δ for each of the variables analysed is available in Table 2.

**Table 2.**
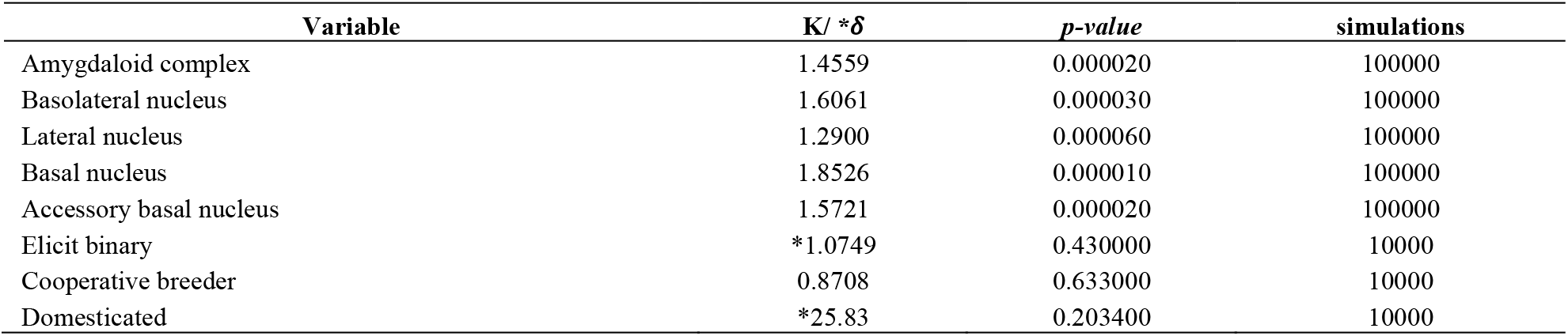
Estimated phylogenetic signal for all the variables used in the analyses. Blomberg’s K was used for continuous variables, while δ was used for discrete ones. Phylogenetic signal is significant for all neuroanatomical variables, and it is very strong (K > 1). Phylogenetic signal is not significant for domestication or cooperative breeding.

Finally, we used the Wheatsheaf Index (Arbuckle et al., 2014) to test for convergent evolution of nuclei volumes in cooperatively breeding and domesticated species. This index allows to explicitly test hypotheses regarding the evolutionary convergence of quantitative traits by incorporating both phenotypic similarities and phylogenetic relatedness. We implemented this using the “test.windex” function in the R package “windex” (Arbuckle & Minter, 2015) which performs a statistical test for convergence. We defined as focal species the ones that are either cooperative breeders or domesticated ones (as per Data S1) and used 10,000 iterations to test for significance.

## 3. Results

Our findings (Figures 1 and 2, Tables 1 and 2) suggest that both domesticated and cooperatively breeding species share adaptations in the amygdala that are fundamentally involved in processing fear-eliciting stimuli (Manassero et al., 2018; Ostroff et al., 2010). First, both cooperative breeding and domestication are associated with a significant increase of the volume of the lateral nucleus relative to BLA (+35% in both cases) and a relative decrease of the basal one (−24% and - 26%, respectively; Figure 1). These results are robust and very similar when regressing L and B against AC: a 46% relative increase of L for both groups, a 25% relative decrease in B for cooperatively breeding species, but no significant effect for domesticated ones. Further, we find that the model with the compound variable “elicit_care” as explanatory variable is the best one (according to the AIC value) in explaining both the increase of the lateral nucleus and the decrease of the basal one. No significant effect is found for the accessory basal nucleus.

**Figure 1.**
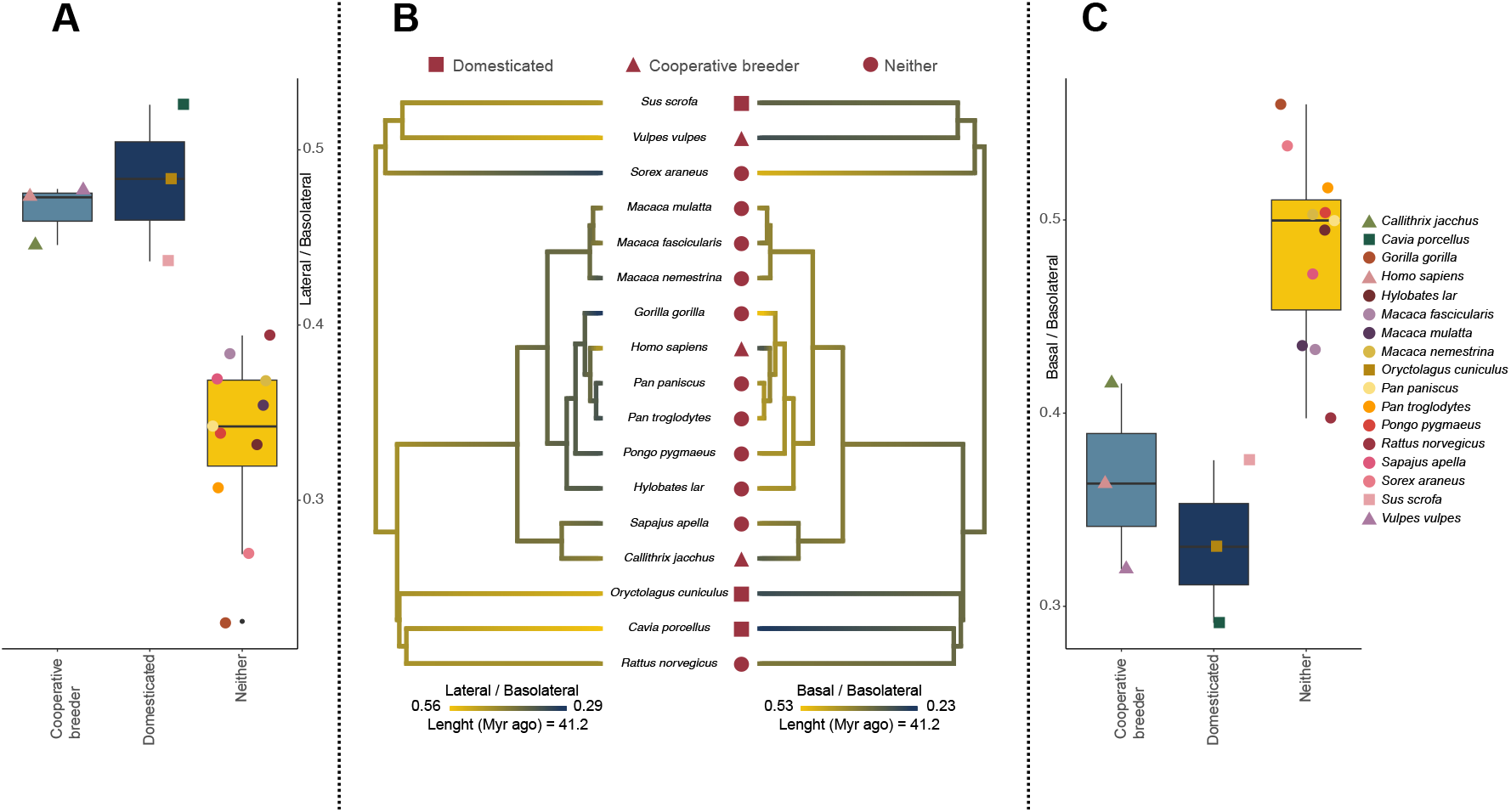
(A) Boxplot indicating the ratio between lateral and basolateral nuclei volumes in cooperative breeders, domesticated species, and others. Each dot represents a species, the legend for the different colors is the same one as reported in panel C. Pairwise corrected p-values of phylogenetic ANOVAs are: 0.003 for *Cooperative breeders* and *Neither*, and 0.018 for *Domesticated* and *neither*. (B) Ancestral state reconstruction of lateral to basolateral volume (left) and basal to basolateral volume (right). In both domesticated species and cooperative breeders, the lateral nucleus is relatively larger (yellow) and the basal one is relatively smaller (blue). (C) Boxplot indicating the ratio between basal and basolateral nuclei volumes in cooperative breeders, domesticated species, and others. Each dot represents a species, identified by a color. Pairwise corrected p-values of phylogenetic ANOVAs are: 0.003 for *Cooperative breeders* and *Neither*, and 0.03 for *Domesticated* and *neither*.

**Figure 2.**
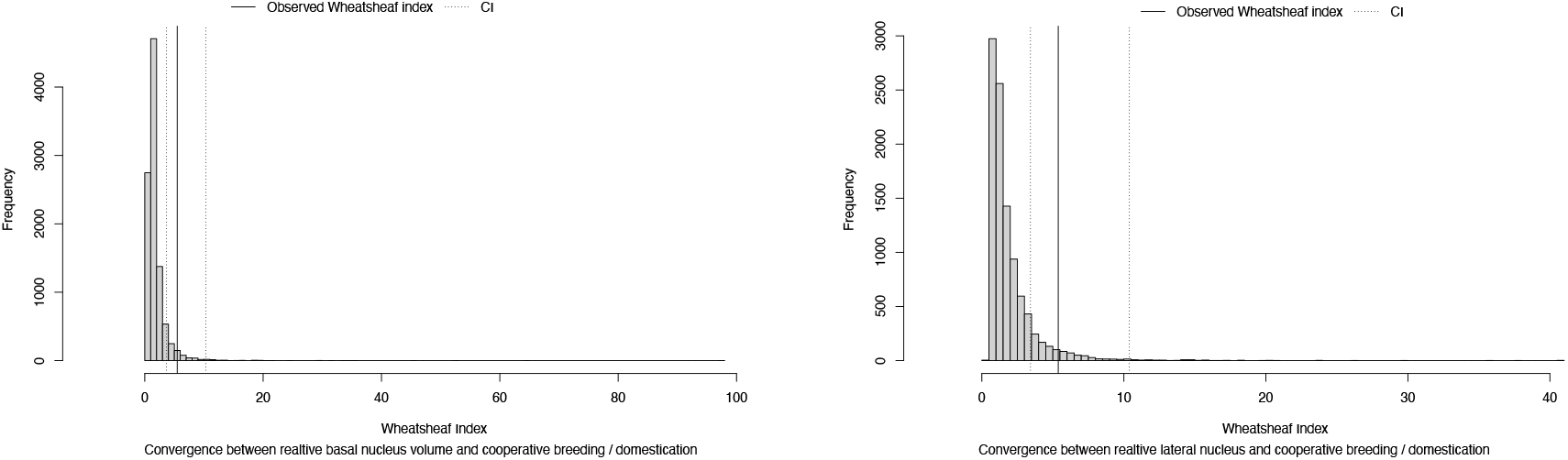
Left: Wheatsheaf Index of the convergence of relative basal nucleus volume in cooperatively breeding and domesticated species (W = 5.49; 95% C.I. = 3.65, 10.31; p = 0.03). Right: Wheatsheaf Index of the convergence of relative lateral nucleus volume in cooperatively breeding and domesticated species (W = 5.38; 95% C.I. = 3.42, 10.49; p = 0.04).

The complete list of all the models tested, their estimates and results are available in Table 1, while the phylogenetic signal for each of the variables used is reported in Table 2.

Second, to test for convergence, we calculate the Wheatsheaf Index (W). The 10’000 iterations support the hypothesis of convergent evolution of cooperative breeding/domestication and: i) relative (to BLA) lateral nucleus volume (W = 5.38; p = 0.04; 95% C.I. = 3.42, 10.49; p = 0.04); ii) relative (to BLA) basal nucleus volume (W = 5.49; p = 0.03; 95% C.I. = 3.65, 10.31; p = 0.03). We thus find robust evidence for a relatively larger lateral nucleus in cooperatively breeding and domesticated species which is the result of convergent evolution.

## 4. Discussion

Our results suggest that reduced reactive aggressiveness, mediated by a relatively larger lateral nucleus, constitutes a shared adaptation in cooperatively breeding and domesticated species. Both processes, domestication and cooperative breeding, apparently tap into the same proximate mechanism: a neurobiological adaptation of the lateral amygdala, likely relating to fear processing and reactive aggressiveness which is adaptative in both cases. For domesticated animals, a reduction of fear towards humans provides adaptive advantages e.g. in terms of food sharing for commensal domestication, as in the case of dogs. Conversely, in cooperatively breeding species the selective pressure for increased social tolerance is entirely different, as it is directed towards conspecifics and confers the adaptive advantage of socially distributing the energetic cost of raising offspring.

Further inquiries with increased samples sizes, regarding the role played by the basal nucleus are warranted by our preliminary results showing that this nucleus (relative to total amygdala volume) is reduced only in cooperatively breeding species and not in domesticated ones. The basal nucleus is implicated, together with the nucleus accumbens, in the avoidance response (Ramirez et al., 2015). A relatively smaller nucleus could potentially correlate with a higher threshold that must be met by a stimulus before eliciting avoidance, thus favoring an approach response. This could be instrumental for mothers to accept that others care for their offspring, and for offspring that individuals other than their mothers care for them.

What are the implications of these findings for human evolution? The human social niche not only required changes in the processing of fearful stimuli and a reduction in fear-induced aggression, but also an increase in prosociality. Whereas most domesticated species are not particularly prosocial, with dogs even loosing this feature during the domestication process (Range & Marshall-Pescini, 2022), empirical evidence shows that cooperative breeding is associated with proactive prosociality (Burkart et al., 2014; Horn et al., 2020). Furthermore, recent research shows that our less reactively aggressive close relatives, bonobos, nonetheless show no prosocility in both prosocial choice task and group service paradigm (Verspeek et al., 2022). A reduction in reactive aggressiveness, potentially mediated by the derived amygdala volumetric patterns found in the present research, is only a prerequisite or stepping stone for the emergence of proactive prosociality. Whereas the presence of the former in humans may be equally linked to self-domestication or cooperative breeding, the latter requires additional selection pressures that are most likely related to cooperative breeding.

## Acknowledgments

We are extremely thankful to Carel van Schaik for providing constrictive feedback and suggestions. We would also like to thank Jeffrey K. Spear for generously sharing his R code, which was used to estimate δ.

## Statements and Declarations

### Ethics approvals

This research did not use samples or materials requiring ethical approval.

### Data accessibility

The author confirms that all data generated or analysed during this study are included in this published article.

### Authors’ Contributions

P.C. conceived the study and carried out data collection and analysis. P.C. and J.M.B. designed the study, interpreted the results, and wrote the manuscript.

